# Reversible Transcriptomic Age Shifts from Physiological Stress in Whole Blood

**DOI:** 10.1101/2024.09.08.611853

**Authors:** Kyungwhan An, Yoonsung Kwon, Jihun Bhak, Hyojung Ryu, Sungwon Jeon, Dougu Nam, Jong Bhak

## Abstract

We developed a genome-wide transcriptomic clock for predicting chronological age using whole blood samples from 463 healthy individuals. Our findings reveal profound age acceleration, up to 24.47 years, under perturbed homeostasis in COVID-19 patients, which reverted to baseline upon recovery. This study demonstrates that the whole blood transcriptome can track reversible changes in biological age induced by stressors in real physiological time, suggesting a potential role for anti-aging interventions in disease management.

## Introduction

The molecular clock estimates biological age beyond chronological age [1]. While DNA methylation has been a primary aging biomarker, its gene regulatory effects remain poorly understood, complicating interpretation [2]. Gene expression offers a more functionally relevant and faster readout than CpG sites [3], reflecting whole-body health conditions in real physiological time [4]. Few studies use a well-established transcriptomic clock for precise measurement of biological age across human diseases. Our study fills this gap by leveraging RNA sequencing to predict chronological age and linking transient pathological states to biological age. We also identify novel aging biomarkers in whole blood, such as *VSIG4*, providing molecular insight into aging.

## Results

### mRNA Accurately Predicts Chronological Age in Healthy Individuals

We trained a machine learning model using RNA-seq data from a cohort of healthy humans and successfully predicted chronological age (Figure 1A-C; Supplementary Figure 1). The predictive model relies on 47 genes associated with lymphoid immune cells (Figure 1D-F).

**Figure 1.**
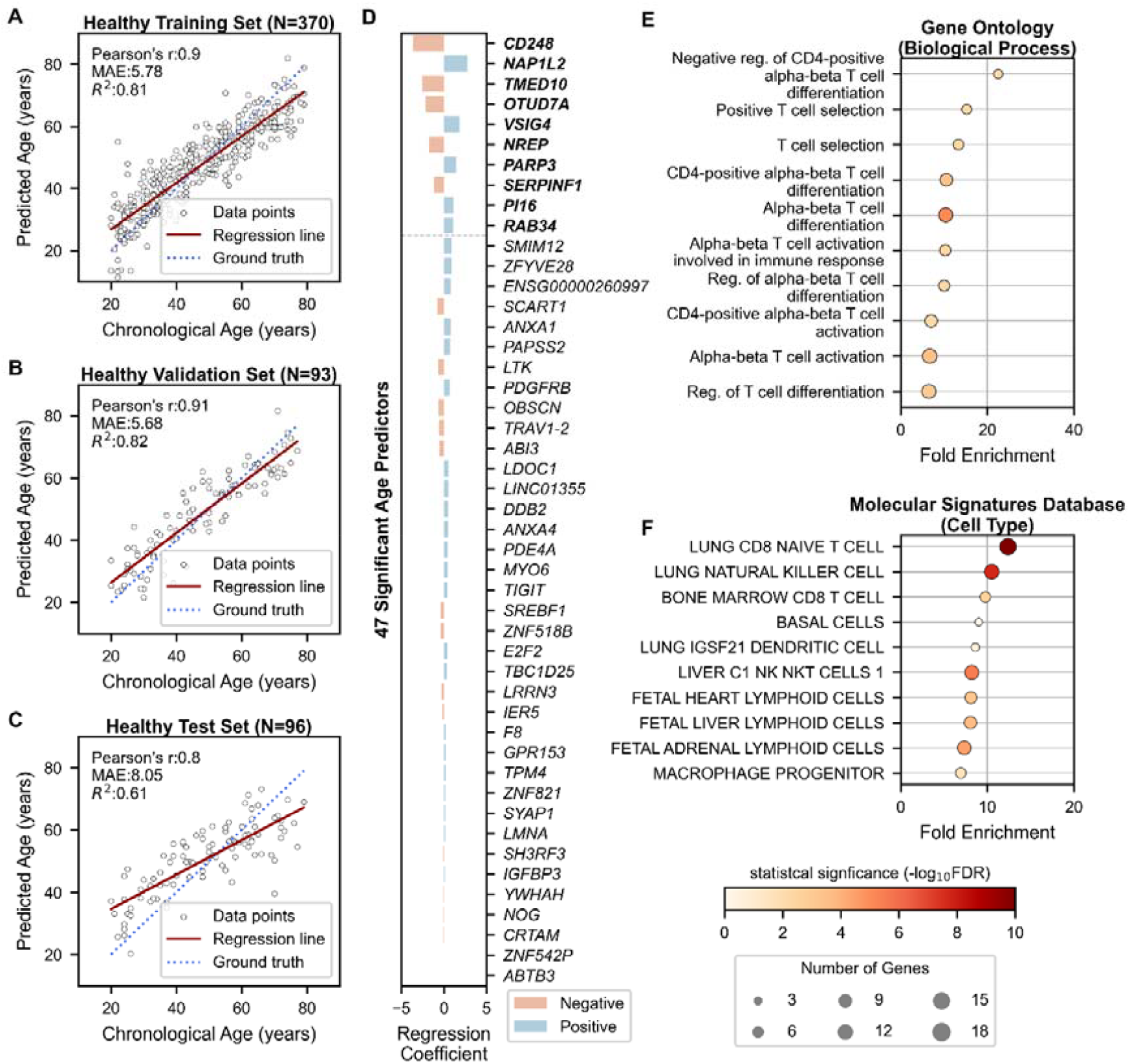
Chronological Age Prediction Using 47 Genes in Healthy Cohorts. **(A-C)** Scatter plots showing the performance of the age prediction model on **(A)** training (N=370), **(B)** validation (N=93), and **(C)** independent test (N=96) data. The x-axis shows chronological age, and the y-axis shows predicted age via the mRNA clock. Each sample is represented by an open black dot, with a solid red line indicating a regression trend and a dotted blue line indicating perfect correlation. **(D)** Bar plot showing genes ranked by their importance in age prediction. The x-axis shows regression coefficients, and the y-axis lists gene symbols of the 47 age predictors. Genes exceeding the threshold are shown in bold, with a threshold being a grey dotted line. Blue and red bars indicate positive and negative associations with aging, respectively. **(E, F)** Dot plots displaying gene-set enrichment results of the 47 age predictors with their 235 co-expressed genes based on **(E)** Gene Ontology (Biological Processes) and **(F)** Molecular Signatures Database (Cell Type). The x-axis represents fold enrichment while the y-axis portrays the top 10 annotated biological functions (FDR < 0.05). Dot color denotes the statistical significance, and size indicates the number of enriched genes. MAE = Mean Absolute Error; r = Pearson’s Correlation Coefficient; R^2^ = Coefficient of Determination; FDR = False Discovery Rate.

We analyzed 12,546 stably expressed genes with age correlation in blood, finding 3,407 genes with no correlation (Supplementary Table 1). These genes are involved in mitochondrial function, metabolic pathways, and chronic diseases (Supplementary Figure 2). We identified 331 genes significantly correlated with age (Supplementary Table 2). From these, we selected 47 genes that collectively predict chronological age with high accuracy (R^2^_train_ = 0.81) using penalized regression (Figure 1A, D; Supplementary Table 3). Model validation across two healthy cohorts confirmed robust performance: 93 in validation (R^2^_validation_ = 0.82) and 96 in a separate test (R^2^_test_ = 0.61) (Figure 1B, C). Samples showed consistent correlation with age across genes (Supplementary Figure 3).

**Figure 2.**
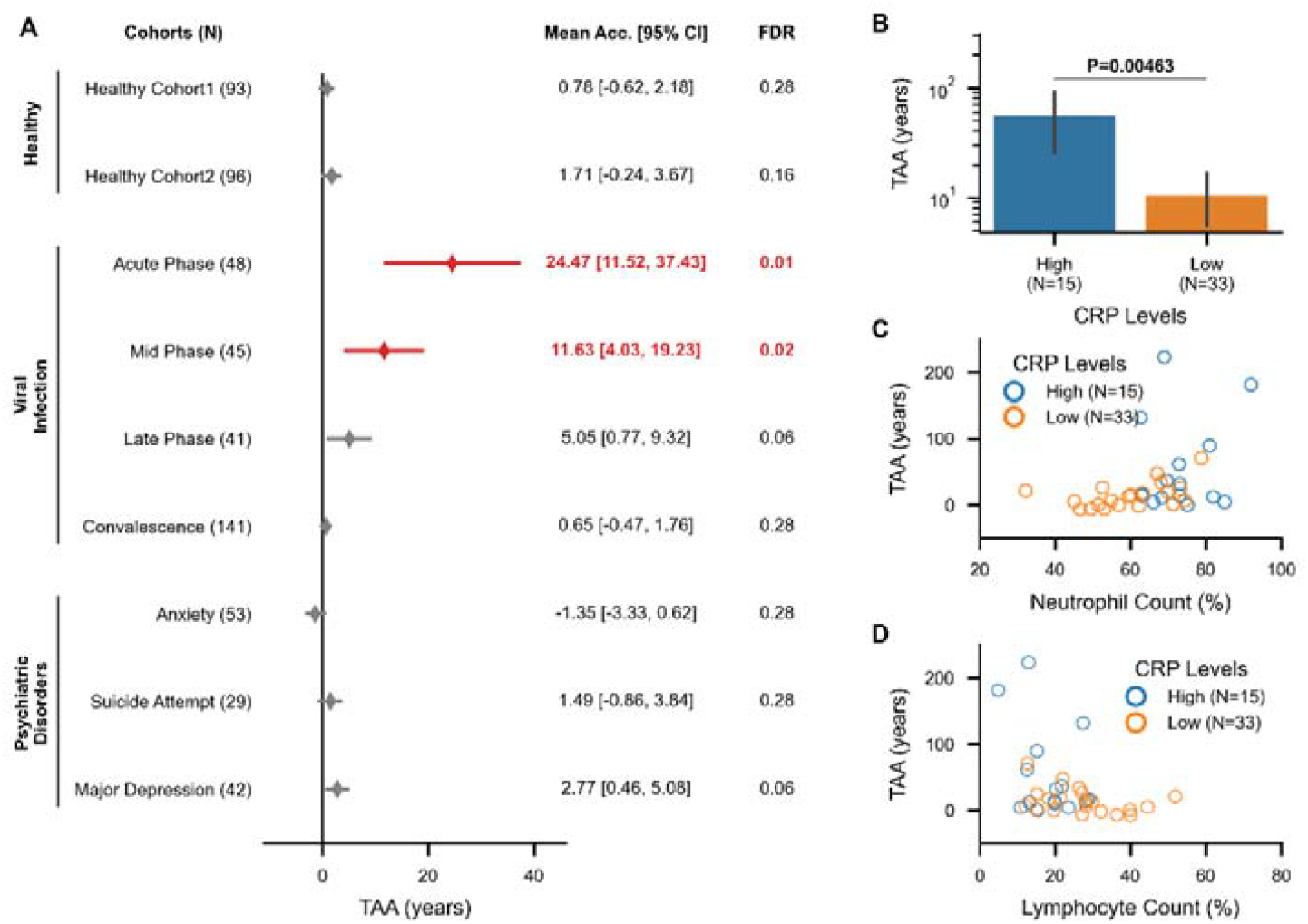
Transcriptomic Age Acceleration (TAA) across Healthy and Diseased Cohorts and Its Clinical Correlates. **(A)** A forest plot illustrates interval estimates of TAA for each phenotype. Red-filled diamonds indicate cohorts with statistically significant TAA (FDR < 0.05), while grey-filled diamonds denote no significance. The x-axis represents TAA in years, and the y-axis lists the study cohorts. Statistics on TAA are displayed on the right side, with bold red figures indicating statistical significance. **(B)** A bar plot shows the difference in mean Transcriptomic Age Acceleration (TAA) between high CRP (>1 mg/dL) and low CRP (≤1 mg/dL) groups among COVID-19 patients at acute phase (N=48) in blue and orange bars, respectively. The x-axis shows the CRP groups, while the y-axis shows the TAA scaled by log10. **(C, D)** Scatter plots depict longitudinal changes in TAA with **(C)** Neutrophil Count (%) and **(D)** Lymphocyte Count (%), categorized by CRP groups. Open dots represent COVID-19 patients. TAA = Transcriptomic Age Acceleration; Mean Acc. = Mean Acceleration; CRP = C-Reactive Protein; N = sample size; CI = Confidence Interval; FDR = False Discovery Rate of the one-sample t-test against μ_TAA_ = 0 (two-sided); P = P-value of Mann-Whitney U test (one-sided).

The first principal component of the 47 age predictors, explaining 21.4% of total variance, effectively stratified our samples by age groups (Supplementary Figure 4A). The 47 age predictors and 235 co-expressed genes were predominantly enriched in T-cell and innate lymphoid cell markers (Figure 1E, F), though not exclusively. For instance, *VSIG4* and *NREP* involve in immunosuppression and blood pressure homeostasis, respectively. These genes have been reported as deleterious signatures of aging across multiple tissues and species [5]. In our model, a unit standard deviation increase of *NREP* level corresponds to a 1.7-year reduction in transcriptomic age, while that of *VSIG4* translates to a 1.8-year increase (Supplementary Table 3). Given the compelling functions, our age predictors are prime surrogate markers with diagnostic implications.

### Transcriptomic Age Acceleration in Response to Physiological Stressors

Disparities in the expression profiles of age predictors were observed among different diseases (Supplementary Figure 4B), despite a near-uniform age distribution (Supplementary Figure 5). Acute viral infection notably increased transcriptomic age by 24.47 years, as measured by the mRNA clock, with levels returning to a physiologically healthy state during convalescence (Figure 2). *VSIG4* emerged as a prominent gene with dramatic changes in expression corresponding to shifts in transcriptomic age (Figure 3). Additionally, transcriptomic age correlated with clinical markers of hyperinflammation (Figure 4).

We applied the model using the 47 genes on two disease cohorts (Supplementary Figure 1) and could not predict the age of some unhealthy individuals (Supplementary Figure 6). We quantified transcriptomic age acceleration (TAA) to find deviations between the chronological and transcriptomic age (Supplementary Figure 7). Our healthy cohorts showed no significant age acceleration (Figure 2). The summarized mean of the two cohorts was no more than 1.25 years (95% CI: -0.62 to 3.67, FDR=0.065) (Supplementary Table 4), confirming the non-diseased status of the healthy samples.

In SARS-CoV-2 infection, longitudinal samples showed a mean age acceleration of 24.47 years during the acute phase (95% CI: 11.52 to 37.43, FDR = 0.01). This declined to 11.63 years (95% CI: 4.03 to 19.23, FDR = 0.02) and 5.05 years (95% CI: 0.77 to 9.32, FDR = 0.06) in the mid and late phases, respectively. Moreover, an independent cohort of 141 convalescent samples showed no evidence of acceleration, with a mean of 0.65 years (95% CI: -0.47 to 1.76, FDR = 0.28) (Figure 2). Expression dynamics of the 47 age-predictive genes among COVID-19 patients support this return to homeostasis, emphasized by dwindling *VSIG4* level (Supplementary Figure 8; Supplementary Table 5). Psychiatric patients exhibited an insignificant mean acceleration overall (0.97 years; 95% CI: -3.32 to 5.08, FDR = 0.29) (Supplementary Table 4).

We found that COVID-19 patients with higher inflammatory status, indicated by C-reactive Protein (CRP), have elevated mean age acceleration (P=0.0046) (Figure 2B). In patients with high CRP levels, the average TAA showed a positive correlation with neutrophil count and a negative correlation with lymphocyte count (Figure 2C, D).

## Discussion

COVID-19 associates with advanced epigenetic age [6]. The virus acts as a pro-aging factor. Virus-induced senescence (VIS) is linked with disease severity through senescence-associated secretory phenotypes (SASPs), causing systemic inflammation [7]. COVID-19 patients show reduced NK and CD8^+^ T cell numbers, loss of proliferative activity, and exhausted senescent phenotypes [8-10]. Our transcriptomic clock aligns with these findings. The marked increase in biological age is accompanied by a state of hyperinflammation. The reduction in acceleration during the decline phase reflects an anti-inflammatory response, including the downregulation of *VSIG4*, a potent T-cell suppressor [11]. This challenges the view of aging as merely a risk factor for infection susceptibility [12]. Senolytics abate complications of viral infections, suggesting therapeutic potential [13]. Our findings also implicate anti-aging interventions in improving vaccine efficacy [14].

Mental health problems are associated with epigenetic age acceleration [15]. Our transcriptomic data could not replicate this observation. Cumulative lifetime, not acute, exposure to psychosocial stress implicates epigenetic age [16]. Disturbances by chronic stressors may be difficult to capture using both DNA methylation and RNA expression.

In summary, our study strengthens the evidence for reversibility of biological age in response to physiological stressors. Transcriptomic biomarkers can monitor the transient changes in biological age. Anti-aging interventions may offer promising strategies to manage biological aging and improve health outcomes.

The set of age predictors here only partially represents the spectrum of genes involved in aging. We focused exclusively on genes with linear age correlations, overlooking non-linear expression patterns [17]. We also neglected mRNA profiles from early and late life stages [18]. Genes outside our scope are biologically important for aging (Supplementary Figure 2).

Expression variability, either due to biochemical or cellular heterogeneity, introduces noise that affects the accuracy and reproducibility of transcriptomic clocks. Very interestingly, stochasticity alone has been shown to predict biological age [19]. Future studies delineating the deterministic and stochastic nature of transcriptomic age are warranted.

## Materials and Methods

### Study Population and Sample Collection

We collected a total of 559 whole blood samples from healthy donors participated in the Korean Genome Project (KGP) [20, 21]. Additionally, we obtained 124 whole blood samples from the Mental Health Cohort [22]. From the COVID-19 Infection and Recovery Cohorts, we collected 146 and 141 whole blood samples, respectively. Out of the 146 COVID-19 Infection Cohort samples, 134 samples were longitudinally collected from 48 subjects over a one-month period, covering the acute (N=48), mid (N=45), and late (N=41) phases of infection (unpublished).

### Bulk mRNA Sequencing using Illumina Sequencers

Whole blood samples collected in PAXgene® Blood RNA Tubes were stored frozen at -80°C. Total RNA extraction utilized the PAXgene Blood RNA Kit from Qiagen following the manufacturer’s protocol. RNA quality was assessed by analyzing 1 μl on the Bioanalyzer system (Agilent) to ensure RNA Integrity Number (RIN) and rRNA ratio met required standards. We used 100 ng of total RNA for library preparation with the TruSeq RNA Library Prep Kit and TruSeq Stranded mRNA Sample Preparation Kit (Eukaryote) for the HiSeq2500 and NovaSeq5000 platforms, respectively, following the manufacturer’s instructions. Library quality was assessed with the Agilent 2100 BioAnalyzer and quantified using the KAPA library quantification kit (Kapa Biosystems). Paired-end (2×101 or 2×151) RNA sequencing was performed on HiSeq2500 and NovaSeq5000 sequencers.

### Bulk mRNA Sequencing using BGI/MGI Sequencers

To enrich polyadenylated mRNA and deplete rRNA, we used the Dynabeads mRNA Purification Kit (Invitrogen). Libraries were assessed for size distribution using the Agilent D1000 ScreenTape. Library preparation was conducted using BGI’s custom protocol or the MGIEasy RNA Directional RNA Library Prep Set (BGI) for the BGISeq500 and DNBSEQ-T7 platforms, respectively, following manufacturer protocols. Library quantification was performed using the Qubit 2.0 Fluorometer with the Qubit DNA HS Assay kit (Thermo Fisher Scientific). Paired-end (2×100 or 2×150) RNA sequencing was conducted on the DNBSEQ-T7RS (MGI) platform.

### Expression Quantification and Processing

Sequenced RNA reads had adapters removed and were filtered for low-quality reads using fastp (version 0.23.1) [23]. The filtered RNA reads were aligned to the human reference genome FASTA (GRCH38 p.13) using STAR (version 2.7.10b) with default settings [24]. Transcripts and respective genes were annotated with their Ensembl ID and gene symbol using the annotation GFF3 file (GENCODE version 43) and RSEM (version 1.3.3) [25]. Raw expression of each gene was estimated by RSEM (version 1.3.3) with default parameters [26]. DESeq2 (version 1.42.0; R package) was used to normalize the expression counts for sequencing depth and RNA library composition [27].

To ensure stable mRNA signals, we excluded non-target genes of mRNA-seq and those with very low expression. Genes with a median expression of zero were removed. We also removed those genes below the grand median of all non-zero expression genes. This reduced the number of input genes from 69,222 to 12,546. The remaining genes had their expression values standardized to Z-scores using “preprocessing.StandardScaler” (scikit-learn version 1.3.2).

### Sample Selection for Training the Age Prediction Model

We randomly selected samples from our RNA-seq dataset to achieve a near-uniform age distribution. Samples from age groups 10s, 80s, and 90s were excluded due to distinct aging mechanisms assumed in these groups. From the initial 463 samples, 370 were assigned to the training dataset and 93 to the validation dataset in an 80:20 ratio using “model_selection.train_test_split” (scikit-learn version 1.3.2). The split was stratified into six age group bins using “np.digitize” (numpy version 1.26.2).

### Finding Age-associated Genes via Simple Correlation Analysis

DESeq2-normalized expression values of each gene were correlated with chronological age using Pearson’s test, restricted to the 370 samples in the training dataset to prevent data leakage. P-values were adjusted for multiple tests using the Benjamini-Hochberg approach with “stats.multitest.fdrcorrection” (statsmodels version 0.14.0). Genes with r > 0.30 and FDR < 0.05 were considered significantly associated with chronological age. A linear regression line was plotted for each gene to observe expression trends across age using “linear_model.LinearRegression” (scikit-learn version 1.3.2). Genes with Pearson’s r < 0.05 and FDR > 0.05 showed no significant age-expression correlation.

### Principal Component Analysis

Principal Component Analysis (PCA) was performed on 47 age-predictive genes to examine distinct expression patterns among samples and detect any systematic biases using “decomposition.PCA” from scikit-learn (version 1.3.2). The analysis used training data from 370 healthy samples to learn principal components. Scores of PC1 and PC2 were plotted in a biplot using matplotlib (version 3.8.2).

### Transcriptomic Age Clock

The LARS (Least Angle Regression) LASSO (Least Absolute Shrinkage and Selection Operator) model was trained on 370 healthy samples the genes of significant age correlation using “linear_model.LassoLarsIC” (scikit-learn version 1.3.2) with default parameters. Given our sample size, we proceeded the feature selection with information criterion (asymptomatically equal to Leave-one-out cross-validation) to prevent over- or under-fitting [28]. For detecting the optimal regularization strength (i.e. alpha), we chose a model with the lowest value of Bayesian information criterion (BIC) by iteratively minimizing the BIC (Supplementary Table 6).

Model validation datasets to test the performance in predicting the biological age. Pearson’s correlation (r), Mean Absolute Error (MAE), and Coefficient of Determination (R^2^) was calculated as measurements for the performance using “pearsonr”, “np.mean”, and “metrics.r2_score”, respectively (scipy.stats version 1.11.4; numpy version 1.26.2 ; scikit-learn version 1.3.2).

### Functional Enrichment of Age-Predictive Genes and their Co-expressed Genes

A gene co-expression matrix was constructed from gene expression data, calculating expression-expression correlations using “pandas.DataFrame.corr” with “pearsonr” option (pandas version 2.1.3). The top five genes highly co-expressed with 47 age-predictive genes were selected for downstream analysis. ShinyGO (version 0.80) [29] was used to functionally annotate genes associated with aging.

### Transcriptional Age Acceleration (TAA)

Transcriptional Age Acceleration (TAA), the difference between predicted (transcriptomic) and chronological age, was calculated for each sample. Prediction error confidence intervals were determined using “sem” (scipy.stats version 1.11.4) and tested for significance using two-tailed, one-sample t-tests with adjustments for multiple comparisons (FDR < 0.05) using “stats.ttest_1samp” and “stats.multitest.fdrcorrection” (statsmodels version 0.14.0).

### Gene Expression Dynamics of Differentially Expressed Genes (DEGs) in COVID-19

Raw read counts estimated from RSEM were compared between 48 COVID-19 subjects longitudinally collected and 370 healthy bloods in the training data at acute (N=48), mid (N=45), and late (N=41) phases. DESeq2 (version 1.42.0; R package) was used to discover differentially expressed genes using Wald’s test. Those genes with baseMean below 10 were removed. COVID-19 significant gene set (i.e., COVID19) was defined as those genes with statistics of |log2FoldChange| ≥ 1 and FDR < 0.05 while the non-significant gene set (i.e., None) as |log2FoldChange| < 1 and FDR > 0.05. The 47 age predictor genes belong to AgePred gene set. Differences in expression levels were tested using Kruskal-Wallis test with post-hoc Dunn’s test for pairwise comparisons, correcting p-values with “bonferroni” option in “posthoc_dunn” (scikit-posthocs version 0.9.0).

### Clinical Correlates of Transcriptomic Age Acceleration (TAA)

Clinical lab values of routine blood tests were correlated with Transcriptomic Age Acceleration (TAA) using “pearsonr” (scipy.stats version 1.11.4). Significance was adjusted for multiple comparisons using “stats.multitest.fdrcorrection” (statsmodels version 0.14.0). We focused on the C-reactive protein (CRP) levels in COVID-19 blood samples during the acute phase. High inflammatory response was defined as CRP > 1mg/dL, and low as CRP ≤ 1mg/dL. The difference in median TAA between the high and low CRP groups was calculated by Mann-Whitney U (MWU) test using “mannwhitenyu” (scipy.stats version 1.11.4). Additionally, we investigated the correlations of TAA with neutrophil and lymphocyte counts (%), stratified by CRP groups.

### Availability of Data and Materials

Both normalized and unnormalized read count matrices used in the analysis can be found in our Github page. Raw sequencing data and materials used in the study are available from the corresponding author upon request. The codes used to generate data and calculate statistics, as well as the respective readme files, are openly available in the Github page: https://github.com/korean-genomics-center/transcriptomic_clock.

## Abbreviations

BIC: Bayesian Information Criterion
COVID-19: COronaVIrus Disease of 2019
FDR: False Discovery Rate
GO: Gene Ontology
KEGG: Kyoto Encyclopedia of Genes and Genomes
KGP: Korean Genome Project
LARS: Least Angle Regression
LASSO: Least Absolute Shrinkage and Selection Operator
MAE: Mean Absolute Error
PCA: Principal Component Analysis
RSEM: RNA-Seq by Expectation-Maximization
SASPs: Senescence Associated Secretory Phenotypes
STAR: Spliced Transcripts Alignment to a Reference
TAA: Transcriptomic Age Acceleration
VIS: Virus-Induced Senescence.

## Author Contributions

K. An, D. Nam, and J. Bhak conceptualized the study. H. Ryu helped data collection. K. An conducted the data analysis. Y. Kwon, H. Ryu, S. Jeon helped the data analysis and interpretation. J. H. Bhak helped with project design. D. Nam helped peer-reviewing the methodology. K. An wrote the manuscript. All authors have contributed to this study and helped improve the paper.

## Acknowledgements

We thank all voluntary participants for donating their blood and the city of Ulsan for supporting the project. We thank Ulsan University Hospital for collecting blood samples of patients with COVID-19, especially Eun-Seok Shin, Ok-Joo Sul, and Seung-Won Ra. We thank Ulsan Medical Center and Korea University Anam Hospital for collecting blood samples of patients with Mental health problems, especially Hyung-Tae Jung. We thank all members of the Korean Genomic Center (KOGIC), especially Changhan Yoon and Youngmin Bhak. We thank the Korea Institute of Science and Technology Information (KISTI) providing us with the Korea Research Environment Open NETwork (KREONET). We also appreciate the Ulsan ICT Promotion Agency (UIPA) which provided us with the BioDataFarm (BDF) system which supports the storage, analysis, and management of the BioBigData.

## Ethical Statements

Our study complies with the ethical guidelines and regulations set forth by the Institutional Review Board (IRB) of the Ulsan National Institute of Science and Technology (UNISTIRB-15-19-A, UNISTIRB-16-13-C, and UNISTIRB-21-15-A), the Ulsan Medical Center (USH.20.013), the Ulsan University Hospital (UUH-2021-04-011-004), and the Korea University Anam Hospital (ED15006).

## Consents

The data in our study are derived from voluntary blood donations, with explicit, comprehensive consent obtained from all participants prior to sample collection. These consent forms clearly articulate the intended use of their data for research purposes and emphasize the voluntary nature of their participation.

