## Supplementary Figures for "Reversible Transcriptomic Age Shifts from Physiological Stress in Whole Blood"

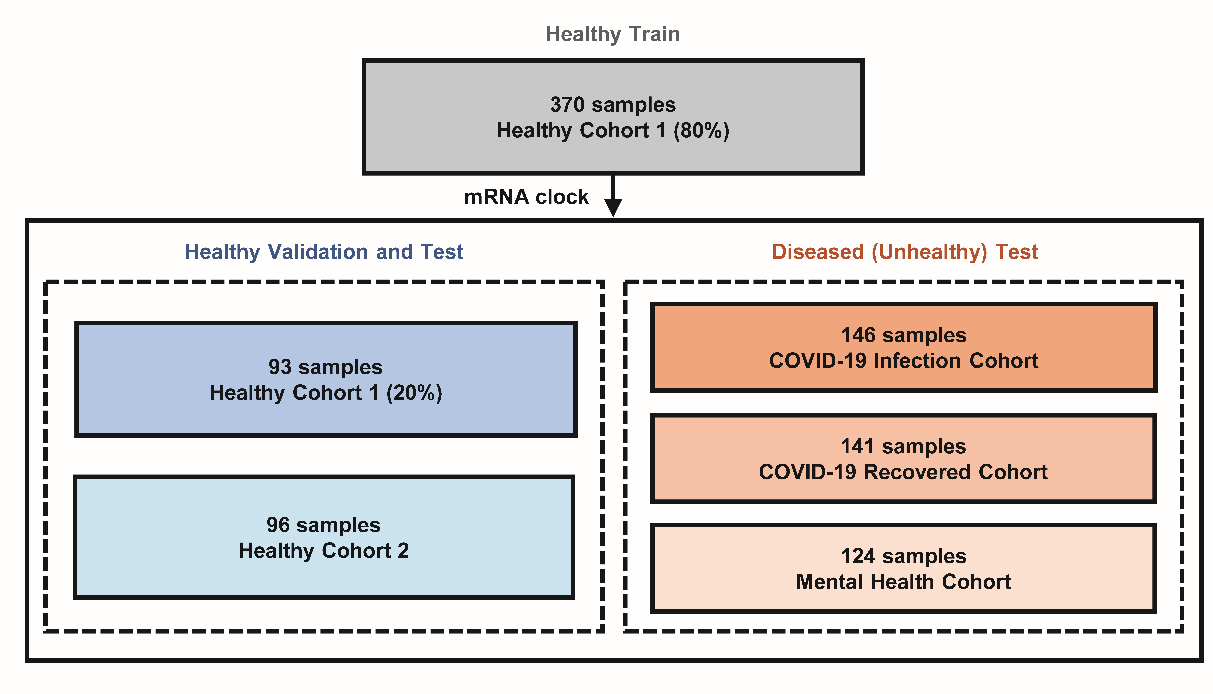

**Supplementary Figure 1. Study Cohorts and Phenotypes.** A total of 970 RNA-seq samples were used. Healthy Cohort 1 for Training (N=370), Healthy Cohort 1 for Validation (N=93), Healthy Cohort 2 for Test (N=96), COVID-19 Infection Cohort (N=146), and COVID-19 Recovered Cohort (N=141), and Mental Health Cohort (N=124).

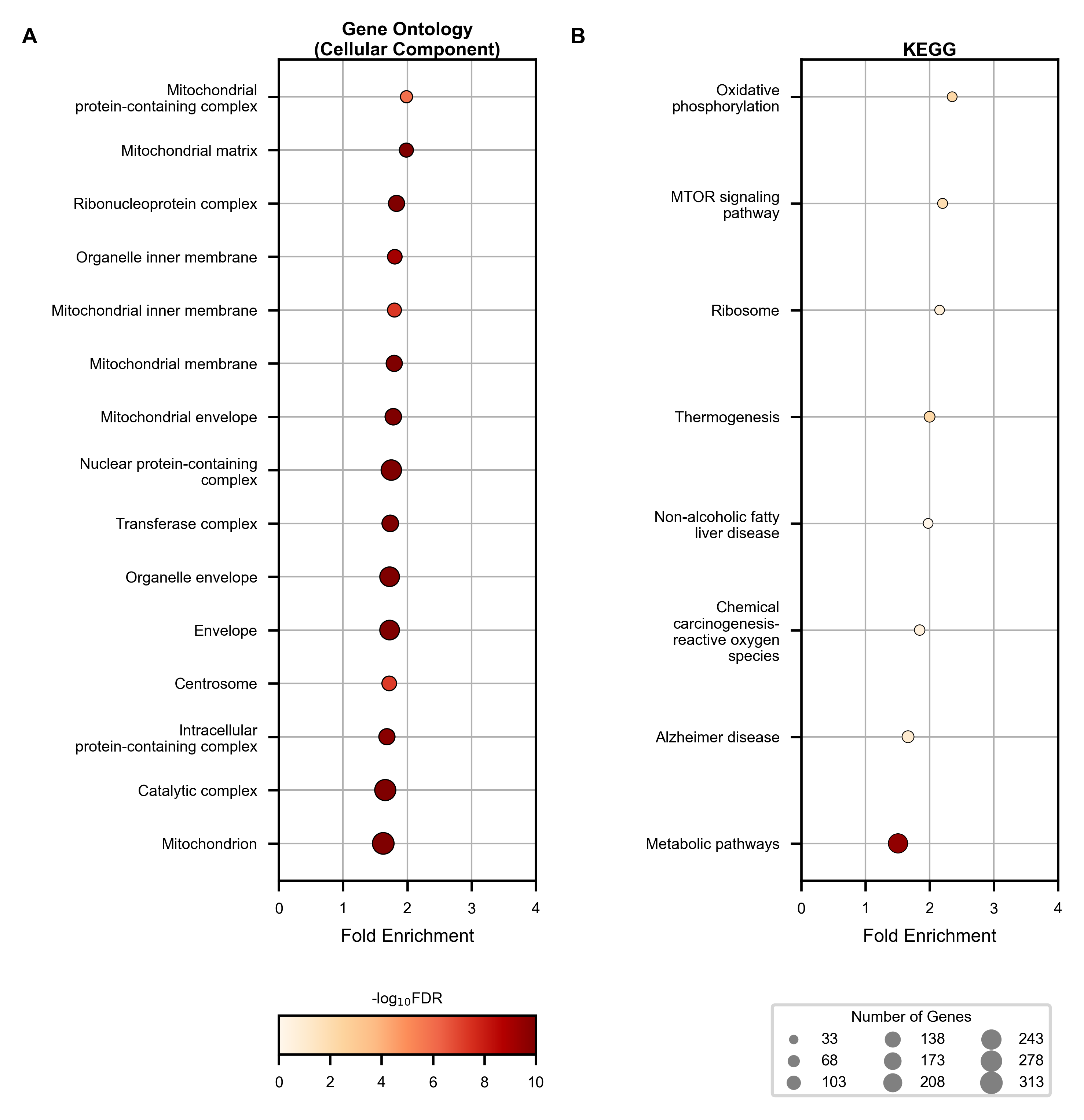

**Supplementary Figure 2. Functional Enrichment of 3,407 Age-Predictive Genes Unassociated with Chronological Age.** Dot plots displaying the biological relevance of the genes according to (**A**) GO Biological Process and (**B**) KEGG. The x-axis represents fold enrichment values while the y-axis portrays the annotated biological functions. The size of the dot represents the number of genes enriched, and the color denotes statistical significance. Black indicates statistical significance, while grey indicates no significance. KEGG = Kyoto Encyclopedia of Genes and Genomes.

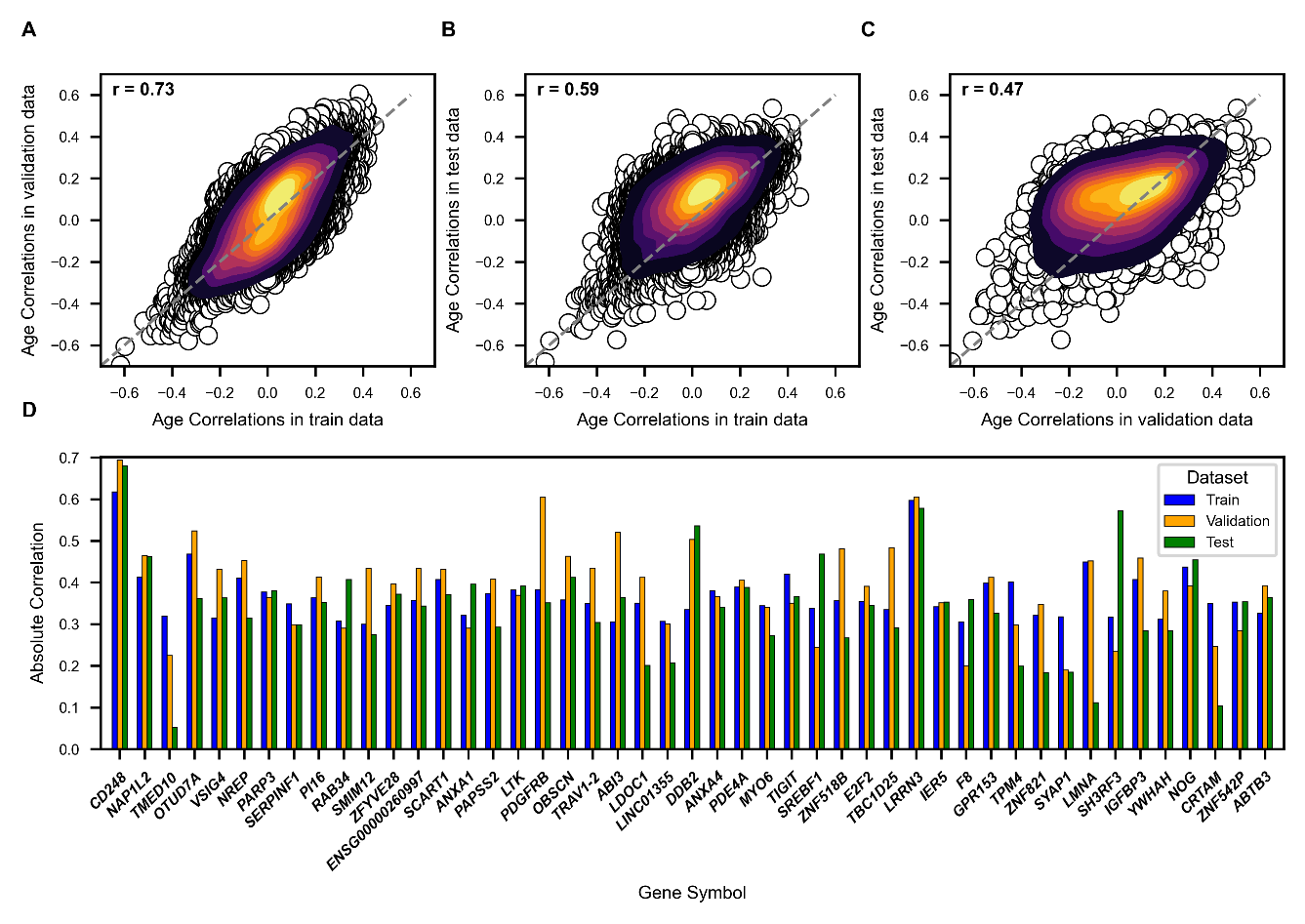

**Supplementary Figure 3. Correlation and Bar Plots Showing Age Correlations of 12,546 Stably Expressed Genes. (A-C)** Scatter plots age showing correlation of 12,546 genes between **(A)** train and validation, **(B)** train and test, **(C)** validation and test data. The x- and y-axes represent Pearson’s r of each gene with chronological age (i.e., Age Correlations). **(d)** Bar plots comparing absolute age correlation of 47 age predictors within healthy cohorts. The x-axis lists gene symbols of the predictors, while the y-axis shows absolute value of Pearson’s r with chronological age.

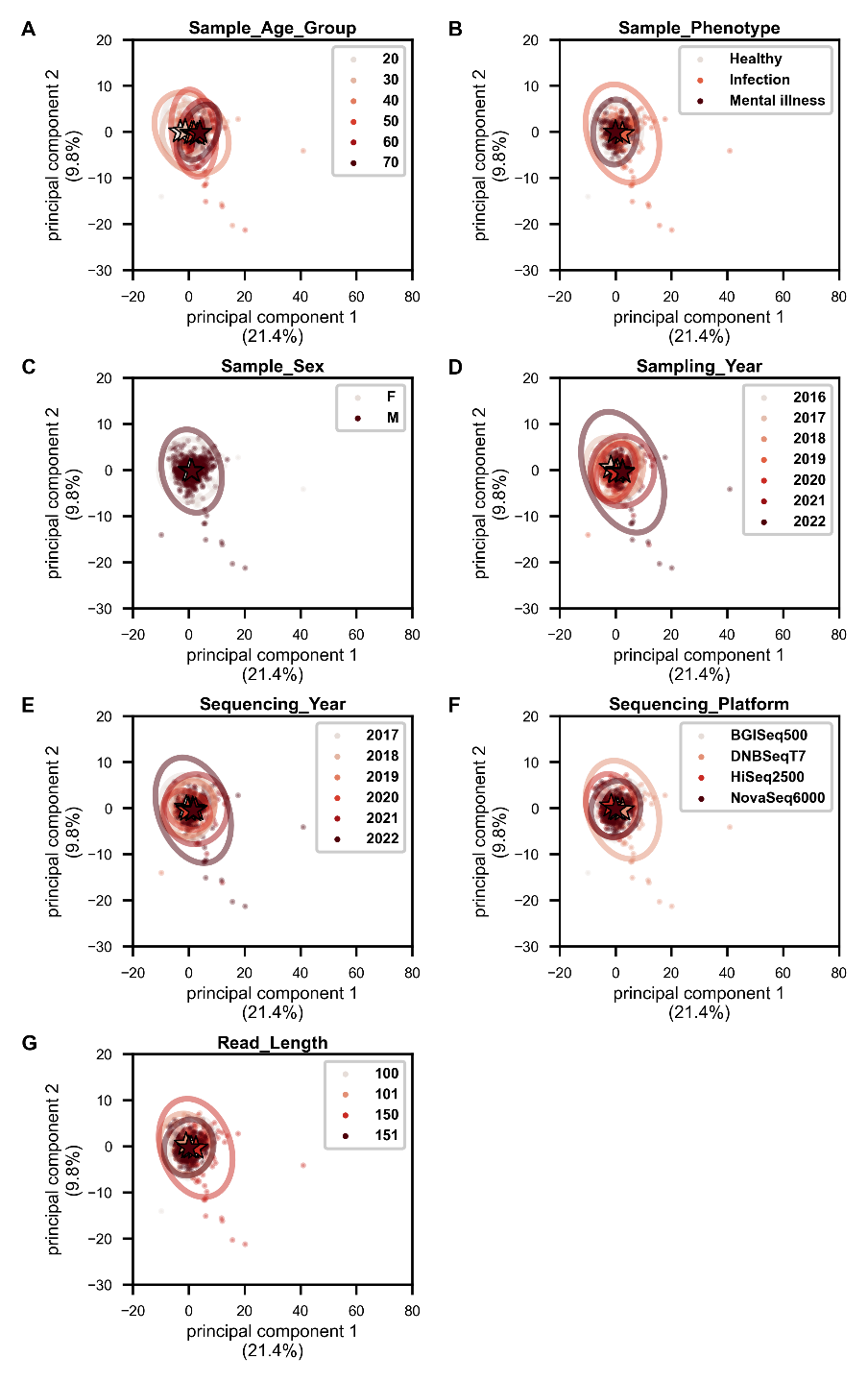

**Supplementary Figure 4. PC Biplots Illustrating the Effect of Disease Phenotypes on Transcriptomic Profiles of 47 Age-Predictive Genes Among 970 Samples.** PC biplots were drawn in relation to **(A-C)** sample information and **(D-G)** batch information, respectively. The x-axis denotes PC1 values of each sample, and the percentage of variance explained. The y-axis displays those of PC2. PC = Principal Component.

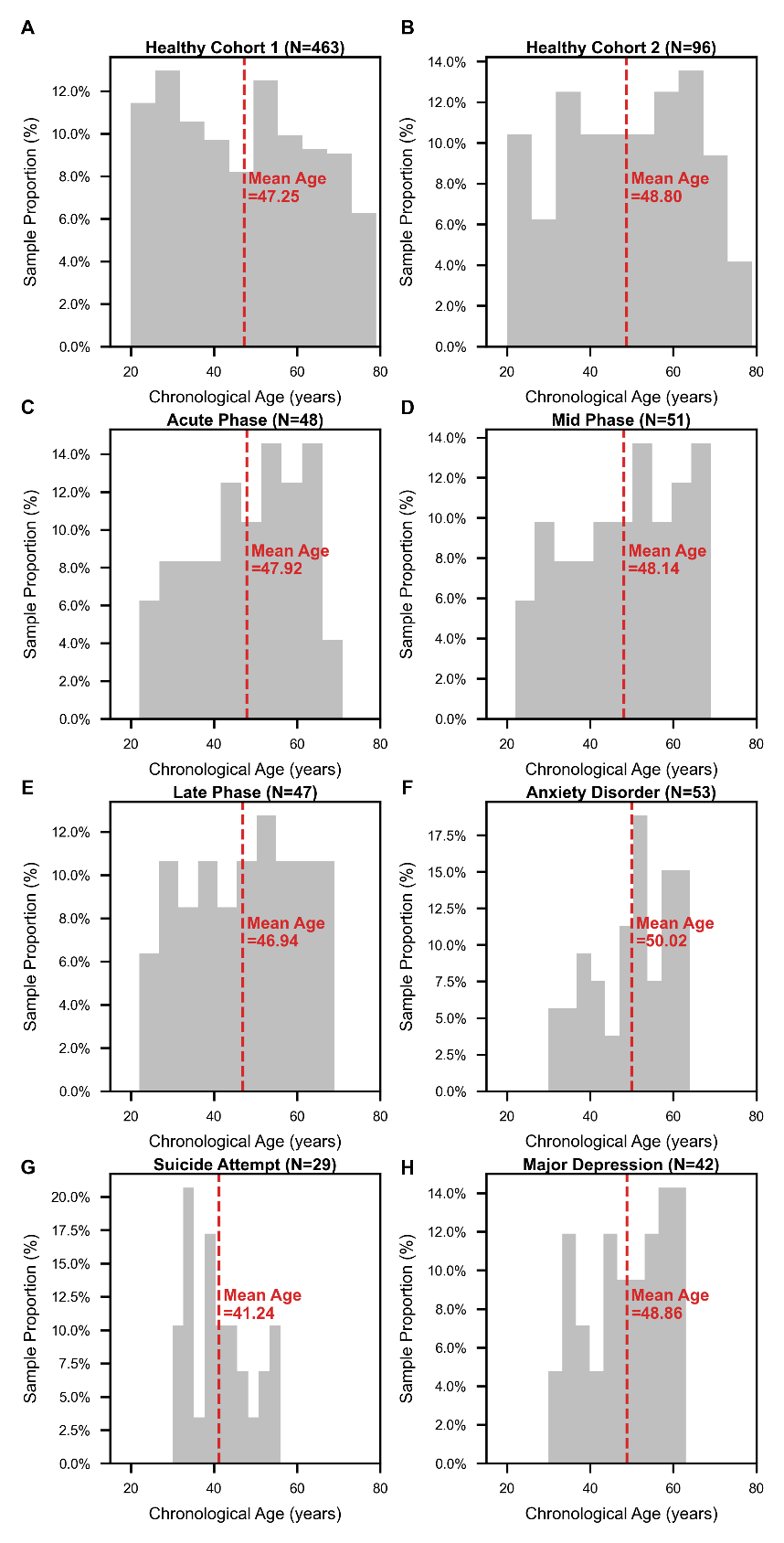

**Supplementary Figure 5. Age Distributions of Study Cohorts.** The x-axis represents the sample age group in years. The y-axis denotes the sample proportion in percentage. A dashed red line and statistics represent the mean age of the study cohorts overall.

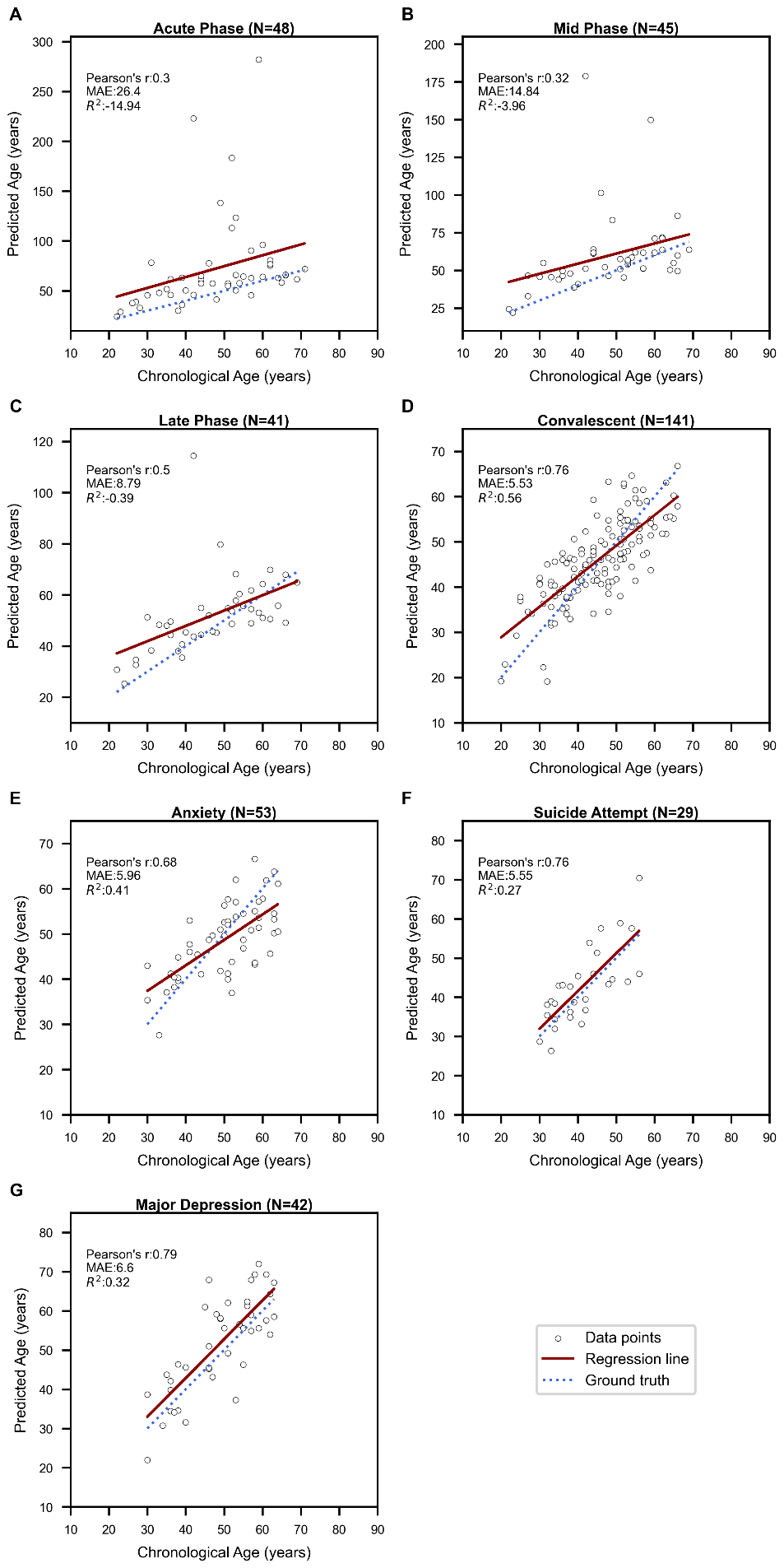

**Supplementary Figure 6. Correlation plots showing the variable prediction accuracies across disease phenotypes.** Scatter plots illustrate the performance of the age prediction model on **(A-D)** COVID-19 patients, and **(E-G)** mentally ill patients. The x-axis corresponds to chronological age, and the y-axis displays predicted age via the mRNA clock. Each open grey dot represents a sample. The dotted line shows perfect correlation, while the solid line represents a linear regression line indicating the general trend of predicted biological age across chronological age. Pearson’s r = Pearson’s Correlation; MAE = Mean Absolute Error; R^2^ = Coefficient of determination.

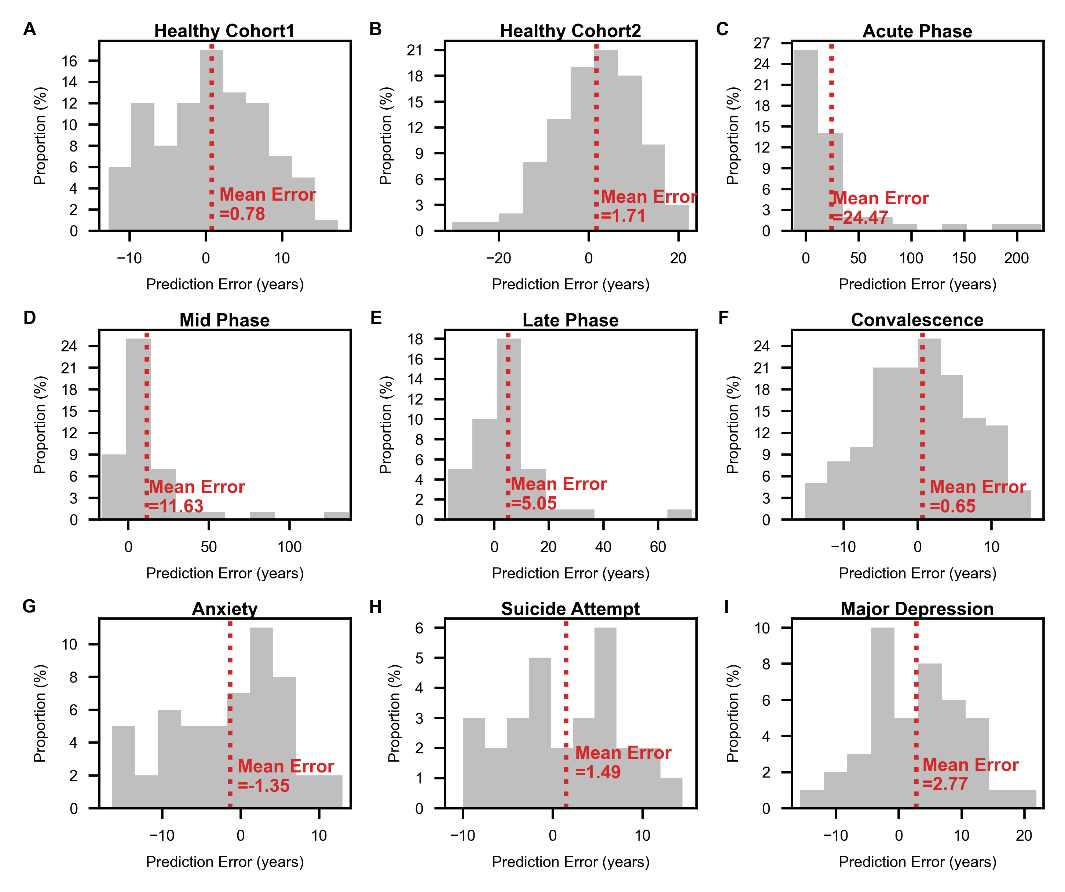

**Supplementary Figure 7. Histograms Showing Distribution of Prediction Errors Across Disease Phenotypes. (A-I)** Grey bars represent the distribution of prediction errors for each study cohort: **(A, B)** Healthy, **(C-F)** COVID-19, and **(G-I)** Mental Health. The dotted red line and annotation denote the mean error for each distribution. The x-axis is the prediction error in years, while the y-axis shows the proportion of samples in percentage.

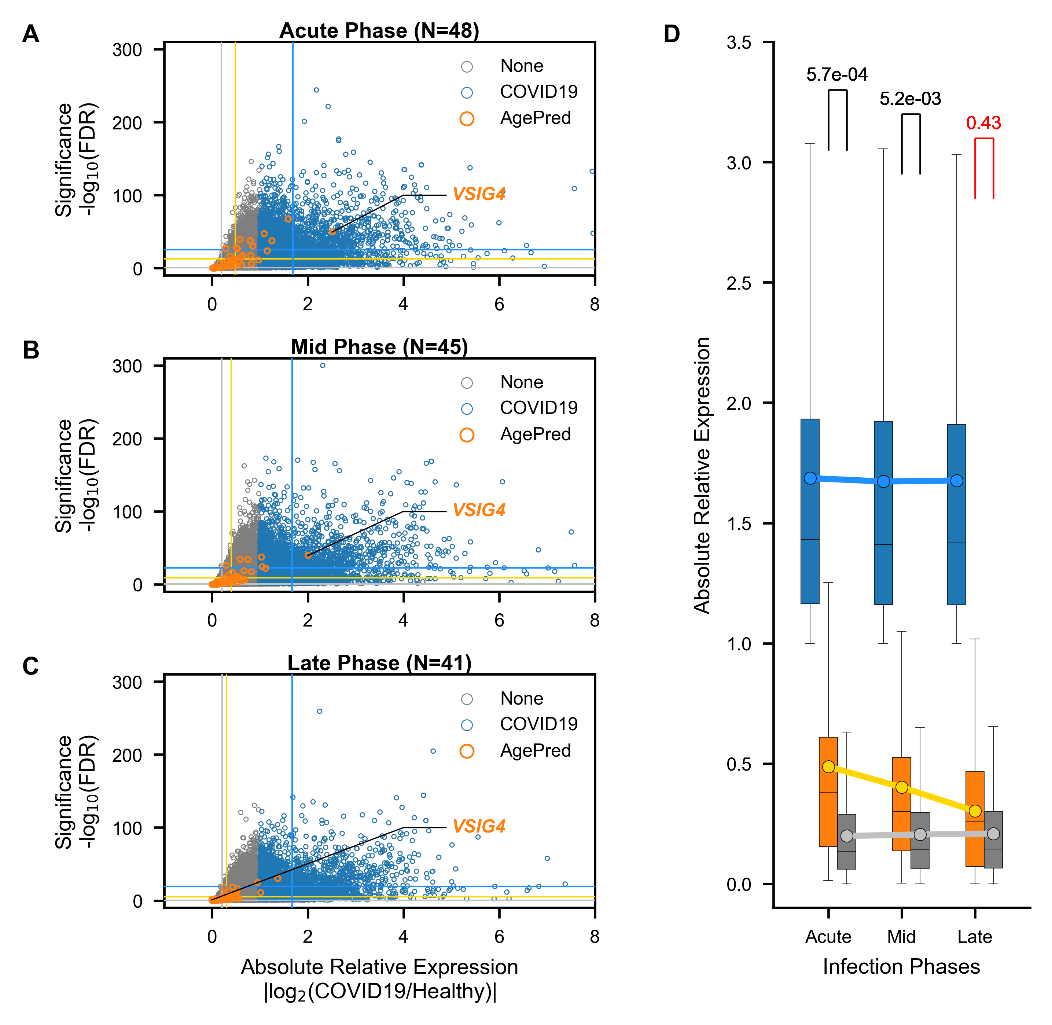

**Supplementary Figure 8. Gene Expression Dynamics of 47 Age-Predictors in COVID-19 Patients. (A-C)** Scatter plots illustrating the relative gene expression in COVID-19 patients at **(A)** acute (N=48), **(B)** mid (N=45), and **(C)** late (N=41) phases, compared to healthy controls. The x-axis denotes absolute relative expression (an absolute log2 value of expression in COVID-19 relative to the healthy), while the y-axis is statistical significance (a negative log10 value of FDR). Each open dot is a gene with blue, orange, and grey colors showing 47 age-predictors (AgePred), COVID-19 significant genes (COVID19; |log2FoldChange| $\geq$ 1 & FDR < 0.05), and non-significant genes (None; |log2FoldChange| < 1 & FDR >0.05), respectively. Solid lines with corresponding colors represent the mean values of each gene set. **(D)** Bar plots comparing the relative gene expression of each gene set across different infection phases. A filled dot represents the mean relative expression in each phase with solid lines portraying the trend of overall relative expression across time. The statistics represent Bonferroni-corrected p-values of post-hoc Dunn’s test between AgePred and None groups. The red figure means no statistical significance.

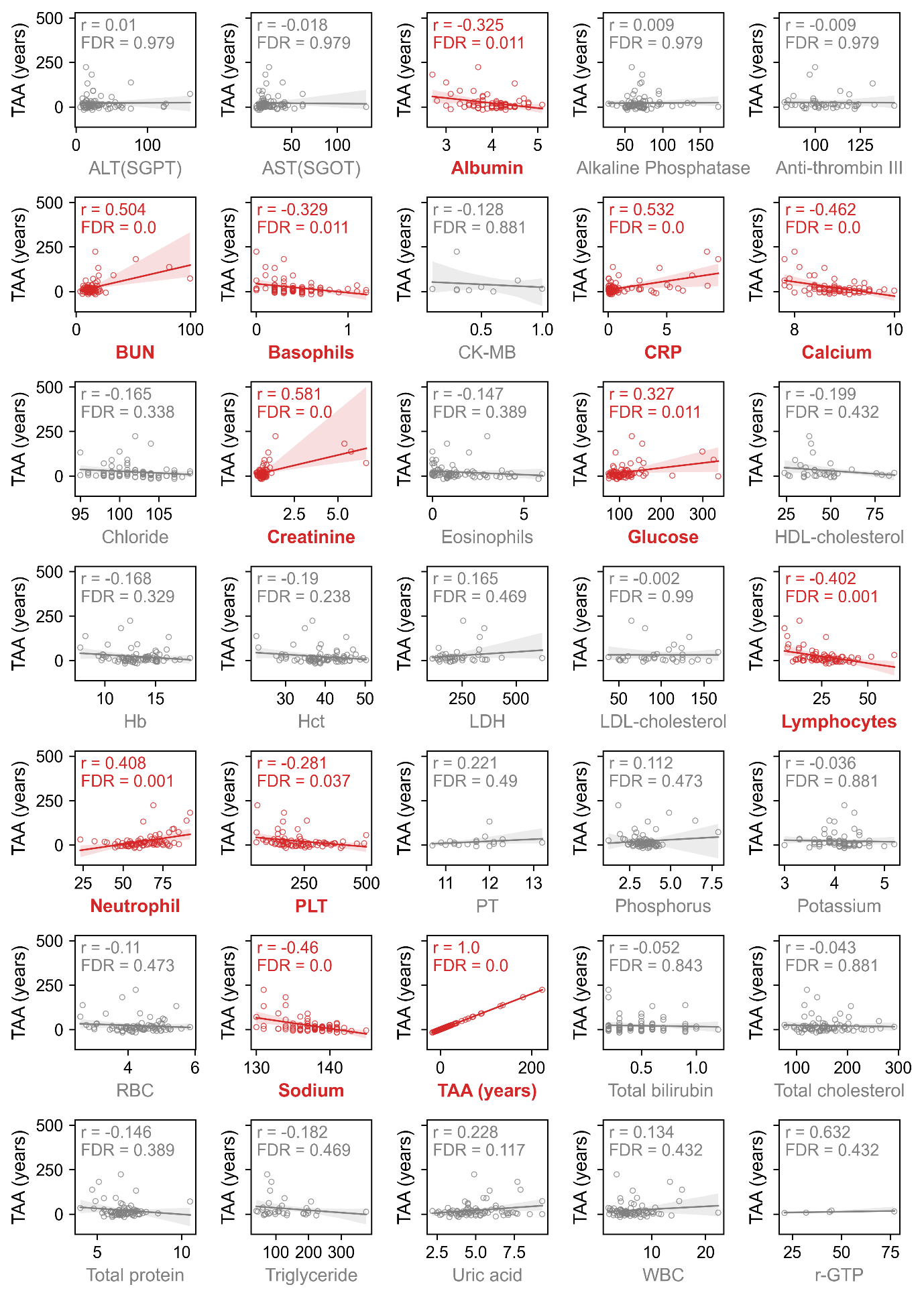

**Supplementary Figure 9. Clinical Correlates of Transcriptomic Age Acceleration (TAA).** Scatter plots show correlations between TAA and routine blood measures (in alphabetical orders) among COVID-19 patients during the acute phase (N=48). Significant associations are denoted in red, while non-significant associations are in grey. Each open circle represents a patient. r = Pearson’s correlation coefficient.
